# Foxes engineer hotspots of wildlife activity on the nutrient-limited Arctic tundra

**DOI:** 10.1101/2021.03.19.436172

**Authors:** Shu-Ting Zhao, Sean M. Johnson-Bice, James D. Roth

**Affiliations:** Department of Biological Sciences, University of Manitoba, Winnipeg, MB, Canada R3T 2N2

**Keywords:** Arctic fox, carrion, ecosystem functioning, nutrient cycling, red fox, scavenging

## Abstract

Predators largely affect ecosystems through trophic interactions, but they also can have indirect effects by altering nutrient dynamics and acting as ecosystem engineers. Arctic foxes (*Vulpes lagopus*) are ecosystem engineers that concentrate nutrients on their dens, creating biogeochemical hotspots with lush vegetation on the nutrient-limited tundra. Red foxes (*V. vulpes*) similarly engineer subarctic environments through their denning behavior, and have recently expanded onto the tundra where they now often occupy historical Arctic fox dens. We evaluated the impact of fox denning activity on the spatial behavior of other tundra wildlife by comparing predator and herbivore visits to 12 natal dens and adjacent control sites over two years using camera traps in northeastern Manitoba, where both fox species are sympatric. Both the capture rates and species richness of wildlife were significantly greater at fox dens relative to control sites. Predators were detected almost exclusively on dens occupied by foxes, where they were observed investigating and scavenging prey remains (carrion, feathers), suggesting carcass presence or fox presence attracts predators to den sites. Caribou (*Rangifer tarandus*) also visited dens more often than control sites, likely attracted by the enhanced vegetation typically found on dens. Our results suggest fox ecosystem engineering affects the spatial distribution of herbivores by enriching vegetation at dens, and other predators by providing carrion. Understanding how predators affect other organisms via non-trophic interactions provides an enriched view of their functional roles within ecosystems.

## 1. Introduction

Predators largely impact other organisms via trophic interactions, but their influence often extends beyond mechanisms such as predation and competition. Predators can affect ecosystem nutrient dynamics by killing prey and redistributing the nutrients and energy across the landscape (Wilmers et al. 2003b, Bump et al. 2009, Schmitz et al. 2010, Monk and Schmitz 2022). For example, brown bears (*Ursus arctos*) consume salmon (*Oncorhynchus* spp.) and transfer marine-derived nutrients to terrestrial habitats, increasing nitrogen concentrations in forest soils (Holtgrieve et al. 2009, Levi et al. 2020). Home sites of predators may often show greater soil and plant nutrient content relative to similar areas due to accumulated nutrients during breeding activities. For instance, the concentration of limiting nutrients and plant species richness are greater around badger (*Meles meles*) setts relative to control areas (Kurek et al. 2014). Predators may also facilitate scavengers and non-scavengers by providing carrion and other prey remains (e.g., feathers for nest-building material) (DeVault et al. 2003, Pereira et al. 2013, Moleón and Sánchez-Zapata 2016, Prugh and Sivy 2020), an often-overlooked trophic interaction that can influence the spatial distribution and diversity of scavenger guilds (Wilmers et al. 2003a, Wilson and Wolkovich 2011, Barton et al. 2013, Moleón et al. 2014).

Organisms that influence the flow of energy and resources by physically modifying their environment are known as ecosystem engineers (Jones et al. 1994). Ecosystem engineers are recognized as important facilitators of community assemblages by generally increasing species richness and diversity in their environments (Jones et al. 1997, Romero et al. 2014, Yeakel et al. 2020). Engineering via soil enrichment and nutrient cycling is particularly important in physically stressful ecosystems where nutrients are often limited, although it is less widely recognized as an engineering mechanism than structural modifications to the environment (Crain and Bertness 2006).

Arctic foxes (*Vulpes lagopus*) are generalist predators and prominent ecosystem engineers in tundra environments (Gharajehdaghipour et al. 2016, Gharajehdaghipour and Roth 2018, Fafard et al. 2020). They are important predators on the tundra that can influence the reproductive success and abundance of shorebirds (Robinson et al. 2014, McKinnon et al. 2014, Flemming et al. 2019), waterfowl (Bêty et al. 2001, Reiter and Andersen 2011, Nolet et al. 2013), and arvicoline rodents (Angerbjörn et al. 1999, Elmhagen et al. 2000, Ims and Fuglei 2005). Arctic foxes also influence tundra environments via non-trophic means through their denning behavior. Den sites are restricted on the tundra, leading Arctic foxes to use established dens for decades or longer (Macpherson 1969, Garrott et al. 1983). They also have the largest litters of any canid, likely a life history adaptation that evolved in response to fluctuating food resources (mainly arvicoline rodents) (Tannerfeldt and Angerbjörn 1996, 1998, Angerbjörn et al. 2004). Nutrient concentrations and vegetative production are therefore greater at dens than other areas on the tundra due to the large amounts of urine and excrement produced on site, repeated site disturbance from digging burrows, and the accumulation of prey remains from adult foxes provisioning pups (Smith et al. 1992, Gharajehdaghipour et al. 2016, Gharajehdaghipour and Roth 2018). By enriching den sites Arctic foxes indirectly alter the biomass and community composition of vegetation at dens (Bruun et al. 2005, Fafard et al. 2020). Ultimately, Arctic fox dens act as ecological hotspots in the nutrient-limited tundra that may influence other wildlife (Gharajehdaghipour and Roth 2018).

The combined effects of climate change and human-derived resource subsidies have facilitated the range expansion of red foxes (*V. vulpes*) onto the tundra (Sokolov et al. 2016, Elmhagen et al. 2017, Gallant et al. 2020). Red foxes are larger than Arctic foxes and can outcompete the latter species for food resources and denning sites (Tannerfeldt et al. 2002, Rodnikova et al. 2011, Gallant et al. 2014, Ims et al. 2017), although the degree to which the two ecologically similar predators compete seems to vary (Gallant et al. 2012, Lai et al 2022). If red foxes do supplant Arctic foxes on the tundra, it is unclear whether they can replicate the functional role of Arctic foxes. However, recent research from temperate and subarctic forests has shown that, similar to Arctic foxes, red foxes can also act as ecosystem engineers by concentrating nutrients at frequently used den sites (Kurek et al. 2014, Kucheravy et al. 2021, Lang et al. 2021, Lang et al. 2022). Thus, tundra dens occupied by red foxes plausibly could have similar ecological impacts as those occupied by Arctic foxes.

To fully understand the functional role of foxes in tundra environments, their non-trophic effects on other organisms must be evaluated. Here, we used camera traps to evaluate how Arctic fox dens influence other wildlife (predators and herbivores) by comparing the frequency animals visited dens versus control sites on the tundra along the western coast of Hudson Bay, in northeast Manitoba, Canada. We hypothesized that fox dens attract wildlife, and predicted other wildlife species would be captured by camera traps at dens more often than adjacent control sites. We hypothesized predators are attracted to prey remains around dens actively occupied by foxes, while herbivores are attracted to the lush vegetation on dens regardless of fox activity. Fox dens in this area were presumably first created and occupied only by Arctic foxes, and the ecosystem engineering influence on dens here have been previously attributed to Arctic foxes (Gharajehdaghipour et al. 2016, Gharajehdaghipour and Roth 2018, Fafard et al. 2020). However, red foxes have been observed using these Arctic fox dens in recent years (J.D. Roth, unpublished data, and this study), and may therefore also be responsible for some of the influence on other wildlife.

## 2. Materials and methods

### 2.1 Study area and study design

Our study occurred in Wapusk National Park, Manitoba, Canada, along the western Hudson Bay coastline, between June 1 and August 10 in 2015 and 2016, during the denning period for foxes in our study area. Fox dens in the area are situated on elevated, well-drained relic beach ridges running parallel to the coast, separated by peat lowlands (Roth 2003). Within this denning area, the density of fox dens is ~17 dens/100 km^2^. These dens are long-lasting, prominent landscape features, with surface areas averaging 563 m^2^ (Gharajehdaghipour and Roth 2018), and previous work has demonstrated the vegetation on dens is enriched relative to other areas along the beach ridges (Gharajehdaghipour et al. 2016, Fafard et al. 2020; Fig. 1, supplementary material Fig. S4). While both Arctic and red foxes have occupied tundra dens in recent years (including in this study), none of these dens were occupied by red foxes during the mid-1990s (Roth 2003). Due to the longevity of fox dens here (at least 10/12 dens from this study were present in the 1980s; Bahr 1989), we therefore attribute almost all of the enhanced vegetation at den sites to Arctic fox ecosystem engineering (Gharajehdaghipour et al. 2016, Gharajehdaghipour and Roth 2018, Fafard et al. 2020). Changes to the biomass and species assemblage of plants on dens is slow on the tundra generally, and on the fox dens specifically, so the recent incursion of red foxes onto the tundra has likely had little impact to date on the vegetation on dens. Red foxes may, however, have immediate impacts on other wildlife through their activities while occupying dens.

**Figure 1.**
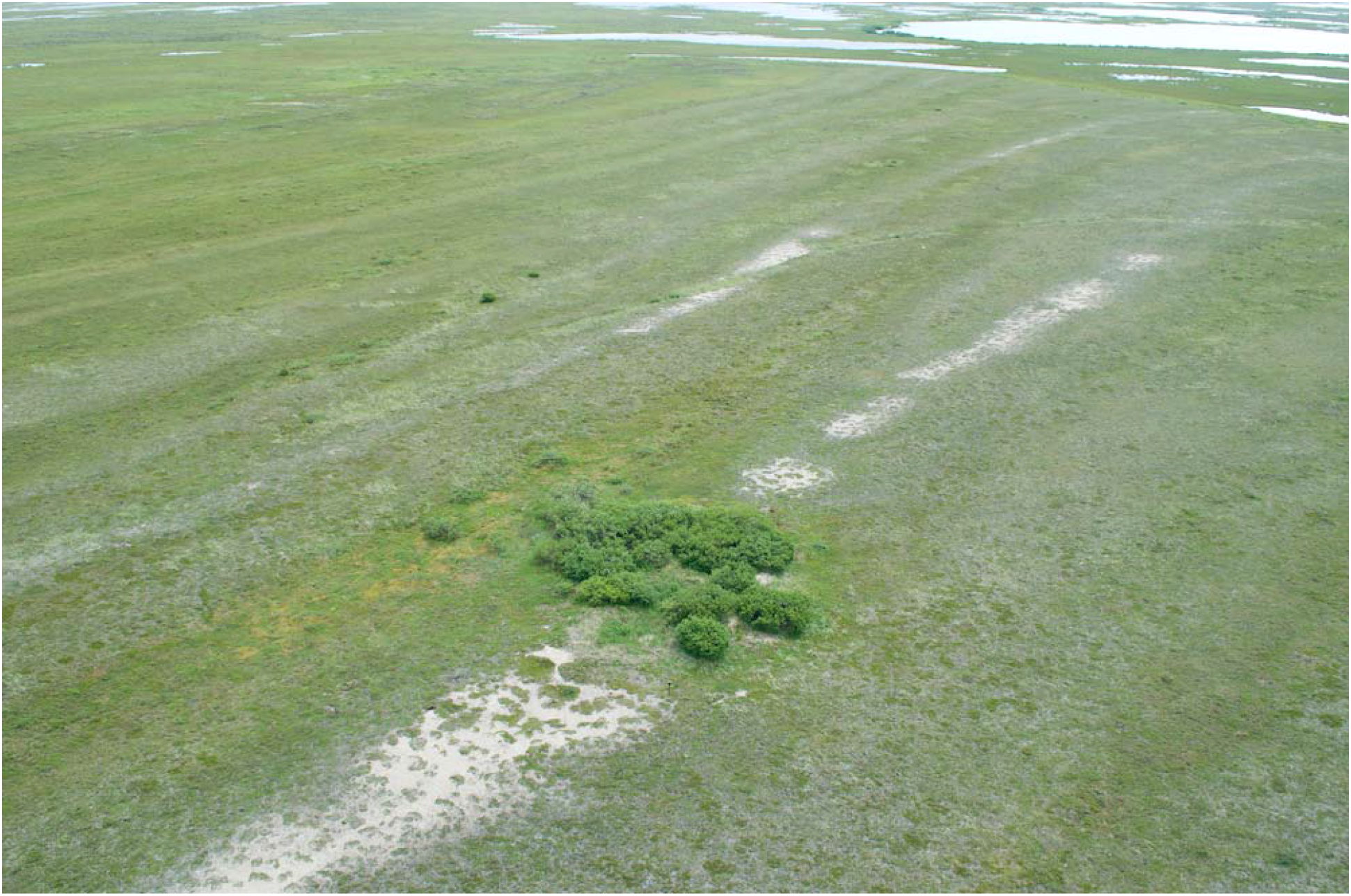
Aerial view of a fox den on the tundra in Wapusk National Park during summer, illustrating the comparably lush, tall vegetation at the den site relative to the surrounding landscape. The elevated beach ridge is also evident, running from the bottom left of the den in the photo up past the top right. We assume that fox dens on the tundra in Wapusk were initially created and occupied by Arctic foxes, and thus attribute most of the distinct vegetative characteristics of dens to Arctic fox ecosystem engineering.

We used a paired study design to compare the frequency that animals were captured on camera traps between dens and control sites, similar to the paired study designs used in previous studies that evaluated the ecological effects of Arctic fox ecosystem engineering at these dens (Gharajehdaghipour et al. 2016, Gharajehdaghipour and Roth 2018, Fafard et al. 2020). We installed cameras onto the same 12 dens each summer (i.e., 2 deployments per den, where deployment refers to times when paired cameras were collecting data). A cased camera was secured on a steel post 1 m above the ground and placed 15 m from the den edge, allowing for maximum visual coverage of the den. We installed a control camera 200 m away from each den, on the same beach ridge, at a similar elevation, slope, and aspect as the den camera. We considered 200 m a sufficient distance such that control cameras would not capture animals on their way to dens.

Three different camera trap models were used in this study (Table S1), but the same camera model was used at each paired site. Cameras were set in motion-detect mode with high sensitivity, and captured a series of three pictures with 10 s interval. If a camera failed mid-deployment or was knocked over by a polar bear (*Ursus maritimus*), we only included data from the time period when both den and control cameras were functioning properly and collecting images. We retained sites that collected data from both cameras for >25 trap nights.

### 2.2 Image evaluation and cataloging behaviors

Animals captured by camera traps were identified to species. We considered events independent if >10 min elapsed between captures (Palmer et al. 2017), unless a different species was captured on camera. With the exception of caribou (*Rangifer tarandus*) herds foraging on den sites, almost all other capture events lasted <10 min. To evaluate whether animals were attracted to dens specifically, we excluded any events where the animal was present <1 min, unless the animal(s) was observed directly interacting with the den or control site (e.g., sniffing the ground, foraging; Fig. 2). The 1-min threshold was used to exclude animals captured on camera that were solely traveling through the camera’s viewshed. We excluded any events if the animal(s) only interacted with the camera (see Supplementary Table S2).

**Figure 2.**
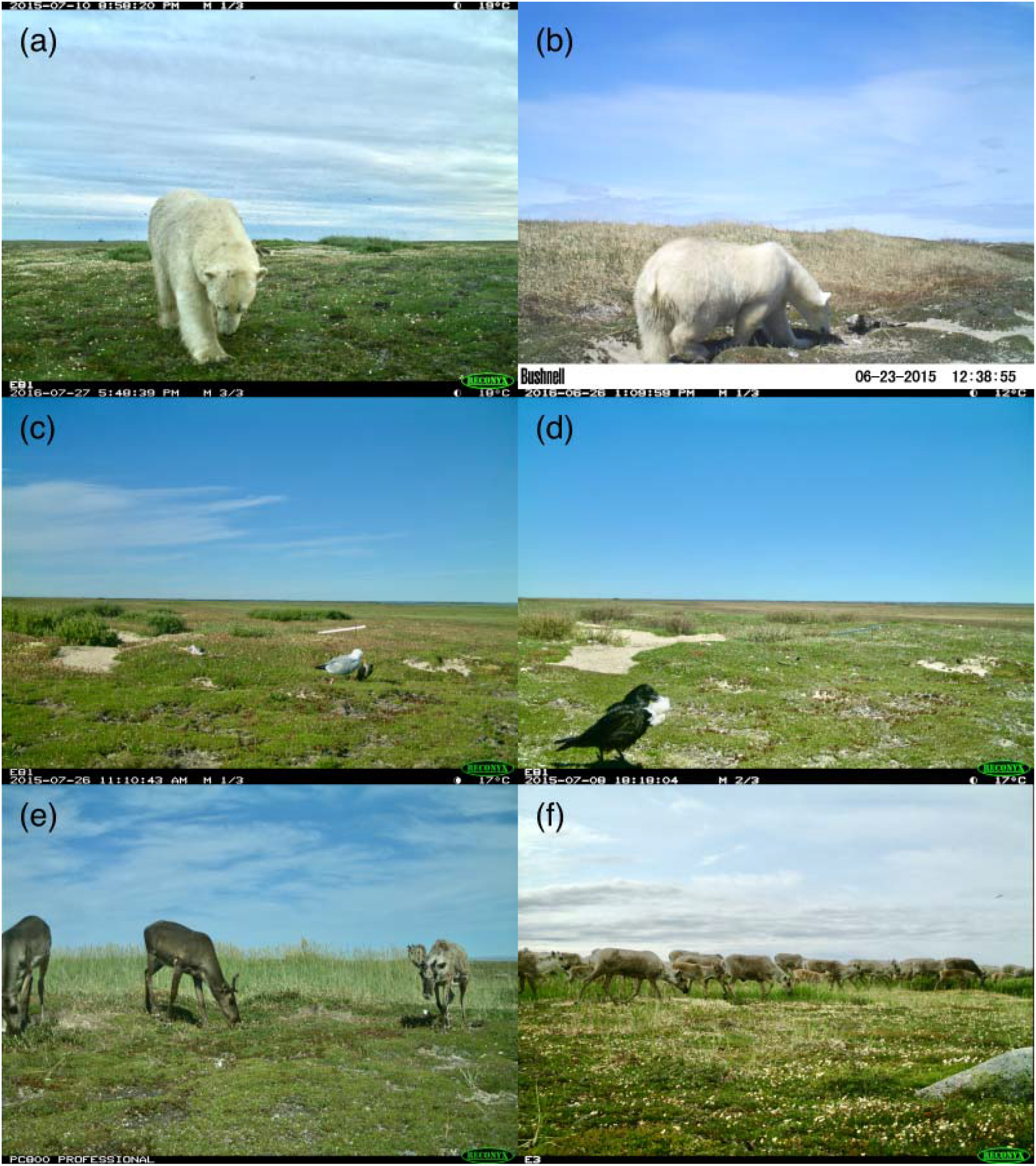
Images captured from cameras placed on dens. Panel (*a*) shows a polar bear (*Ursus maritimus*) leaving a den after interacting with an adult red fox (*Vulpes vulpes*) (tail and hind leg are visible behind the bear). Panel (*b*) shows a polar bear sniffing/investigating a goose carcass. Panels (*c,d*) show a herring gull (*Larus argentatus*) and a raven (*Corvus corax*) removing prey remains/feathers from a den. Panels (*e,f*) show groups of caribou (*Rangifer tarandus*) foraging at fox dens.

For each event, we counted the maximum number of individuals observed within a single image. Only animals that were observed directly on the den or in front of the control camera (i.e., <30 m from the cameras) were included in our counts. Animals captured only off the den or in the background of images were not counted.

We determined whether dens were ‘active’ or ‘inactive’ based on the frequency that foxes were captured on camera. Dens where foxes were photographed on the den at least once per ten trap nights were considered ‘active’.

### 2.3 Statistical analysis

We evaluated our hypothesis that wildlife are attracted to fox dens by comparing capture rates at dens and control sites using generalized linear mixed-effects models implemented in the ‘glmmTMB’ package (Brooks et al. 2017) in R (R Core Team 2020). We compared (1) species richness, (2) number of all wildlife captures, (3) number of predator captures, and (4) number of herbivore captures between the paired den and control sites. To evaluate our hypothesis that predators are attracted to dens by fox activity while herbivores are unaffected, we compared (5) species richness, (6) number of predator captures, and (7) number of herbivore captures between active and inactive dens. Each model was fit with either a negative binomial or Poisson distribution. We assessed model residuals using simulation diagnostic methods from the ‘DHARMa’ package (Hartig 2020), and adjusted the distribution family only if model convergence or overdispersion issues occurred.

For each fitted model, the number of unique species or wildlife (i.e., total number of animals, not the number of events) captured during each deployment was the response variable, while year and treatment (den/control) or activity status (active/inactive) were fixed covariates. Den ID was input as a random effect for all models. Although different camera trap models were used in this study, paired den/control sites had the same camera model to account for detection biases. Exploratory analyses revealed that including camera model as a random effect term had no influence on any parameter value estimates in our models, so we elected not to include that term for simplicity. To control for variable sampling duration, we used the log of trap nights as an offset term in each model. We obtained coefficient estimates, standard errors, confidence intervals, Z-values, and *p*-values for each parameter using the ‘summary’ and ‘confint’ functions in R.

## 3. Results

### 3.1 Comparing wildlife visits between dens and control sites

We retained data from 16 paired den-control deployments (8 of 12 dens per year) to evaluate the influence of fox activity on wildlife capture rates. Among the 16 paired deployments, four dens collected sufficient data from both survey years (8 deployments), while the other eight dens collected sufficient data from both cameras for only one year (8 deployments). We detected four species (other than foxes) on control cameras, compared to 11 species detected on den cameras (Table 1).

**Table 1.** The number of captures by species on dens and paired control sites.

| Species | Category | Captures on control cameras | Captures on den cameras<sup>1</sup> |
| --- | --- | --- | --- |
| Polar bear (*Ursus maritimus*) | Predator | 0 | 7 |
| Wolf (*Canis lupus*) | Predator | 0 | 6 |
| Grizzly bear (*Ursus arctos*) | Predator | 0 | 1 |
| Common raven (*Corvus corax*) | Predator | 0 | 13 |
| Sandhill crane (*Grus canadensis*) | Predator | 0 | 7 |
| <sup>2</sup>Eagle (*Aquila chrysaetos* or *Haliaeetus eucocephalus*) | Predator | 1 | 2 |
| Snowy owl (*Bubo scandiacus*) | Predator | 0 | 1 |
| Herring gull (*Larus argentatus*) | Predator | 0 | 1 |
| Caribou (*Rangiferus tarandus*) | Herbivore | 109 | 269 |
| Canada goose (*Branta canadensis*) | Herbivore | 67 | 115 |
| Snow goose (*Chen caerulescens*) | Herbivore | 25 | 140 |

<sup>1</sup>Two Arctic hares (*Lepus arcticus*) were also detected on a den camera trap whose paired control camera failed.
<sup>2</sup>Observations of eagle species were lumped because it was difficult to distinguish between juvenile golden eagles (*Aquila chrysaetos*) and bald eagles (*Haliaeetus eucocephalus*) from photographs.

Compared to control sites, dens were visited by more wildlife species (i.e., greater total species richness; β_den_ = 0.647, 95% confidence interval [CI]: [0.131, 1.162]), and a greater number of wildlife were likewise detected at dens (β_den_ = 0.994, 95% CI: [0.350, 1.638]) (Fig. 3a, b; Supplementary Table S3). Dens were visited more often by both predators (β_den_ = 3.638, 95% CI: [1.652, 5.623]) and herbivores (β_den_ = 0.83, 95% CI: [0.146, 1.515]) than control sites (Fig. 3c, d). However, a post-hoc analysis revealed that the difference in herbivore visits was driven by caribou (β_den_ = 0.850, 95% CI: [0.131, 1.569]), as goose visits did not differ between den and control sites (β_den_ = 0.743, 95% CI: [−1.861, 3.346]); Supplementary Table S3).

**Figure 3.**
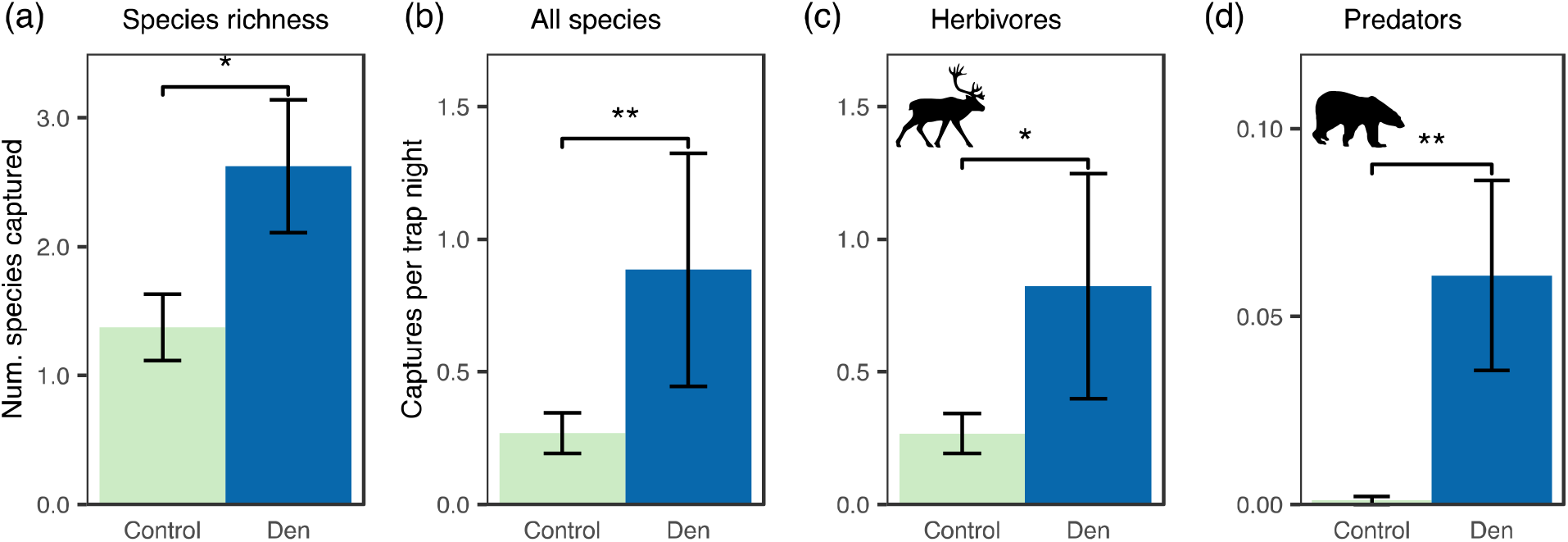
Species richness (*a*) and average daily capture rates (number of wildlife/trap night) of (*b*) all wildlife species, (*c*) herbivore species, and (*d*) predator species at fox dens and paired control sites (\**p*<0.05; \*\**p*<0.01).

### 3.2 Comparing wildlife visits between active and inactive dens

We used data from 18 den camera deployments (10 from 2015, 8 from 2016) to compare species richness and wildlife capture rates between active and inactive dens (the two additional dens from 2015 had control cameras that did not collect sufficient data). Based on the frequency that foxes were detected on cameras, 6/18 dens were determined to be ‘active’. Three of the six active dens were occupied by red foxes, and only three of the active dens produced pups (two Arctic fox dens, one red fox den); the other three active dens had adult foxes using the dens regularly but failed to produce pups. Despite no pups at these other three active dens, adult foxes were captured on camera bringing back prey remains to the dens. Only one den was active both years, used by red fox adult(s) in 2015 and Arctic fox adults and pups in 2016.

Active dens were visited by more wildlife species (i.e., greater species richness) than inactive dens (β_inactive_ = −0.718, 95% CI: [−1.139, −0.296]). The total number of predators observed was greater at active dens (β_inactive_ = −2.556, 95% CI: [−3.584, −1.527]) (Fig. 4a, b), while herbivore visits did not differ between active and inactive dens (β_inactive_ = −0.474, 95% CI: [−1.762, 0.815]). The random effect of ‘den ID’ was not influential for all den activity models except for the model comparing the total number of predators between active and inactive dens.

**Figure 4.**
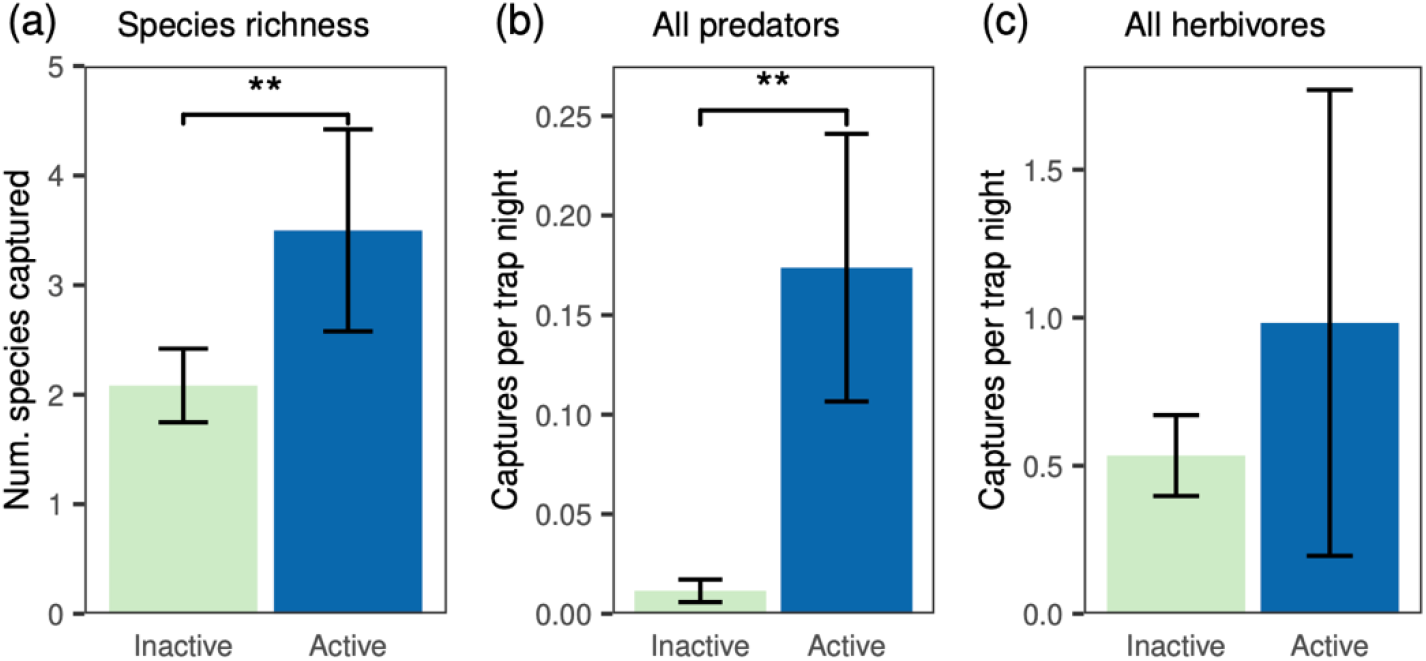
Total species richness (*a*) and average daily capture rates (number of wildlife/trap night) of (*b*) all predators and (*c*) all herbivores between active and inactive fox dens (\*\**p*<0.01).

## 4. Discussion

We observed significantly greater species richness and greater capture rates of wildlife at fox dens than adjacent control sites (Fig. 3a, b), suggesting Arctic foxes create hotspots of wildlife activity on the tundra through their ecosystem engineering. Even with our relatively low sampling effort compared to some camera trap studies, fox dens were found to be visited more frequently by wildlife compared to control areas (e.g., 38 predators detected on dens vs. 1 on control cameras over the same sampling period; Table 1). These differences may be conservative, as several dens had tall vegetation that affected the visibility range of the camera (e.g., Fig. 2e), so we may not have detected all wildlife at den sites since the vegetation may have obstructed some animals. The size of fox dens also varied considerably, and some camera deployments were unable to capture the entire den within its viewshed. Consequently, our detection data across dens may be biased low compared to control sites that were unobstructed by vegetation. Thus, the ecosystem engineering effects of Arctic foxes, and likely red foxes, appear to extend beyond physical modifications of the environment by influencing the spatial distribution of other tundra wildlife by attracting them to dens. These other wildlife likely excrete waste during their visits to dens as well (e.g., Supplementary Fig. S1), creating a positive feedback that may further enhance nutrient enrichment at den sites.

Predators were captured almost exclusively on dens actively occupied by foxes (Fig. 3d, 4b-d, Table 1), which appears to drive the difference in species richness between den/control sites, and active/inactive dens (Fig. 3a, 4a). Prey remains are often littered around active fox dens during the breeding season (Garrott et al. 1983, Roth 2003), and numerous predators were captured on camera scavenging or otherwise investigating prey remains (Fig. 2b-d). An alternative interpretation of these observations may be that the avian predators were removing feathers from these carcasses for nest building materials, as has been observed in other areas (Moleón and Sánchez-Zapata 2016). Given the observations from this study and elsewhere in the Arctic (Mallory 1987) of predators scavenging and removing prey remains from fox dens, we suggest foxes facilitate predators by provisioning carrion and other prey remains (e.g., feathers) at a concentrated location. Since fox dens are such long-lasting, prominent features on the landscape, predators may learn to associate dens with carcass availability and routinely check them for supplemental resources during summer.

Most of the predators we detected in this study have been documented to kill fox pups (e.g., Garrott and Eberhardt 1982, Chevallier et al. 2015), so it is also possible that predators may visit fox dens for hunting opportunities. For instance, predation by eagles (*Aquila chrysaetos* and *Haliaeetus eucocephalus*) may be a substantial source of pup mortality in some areas (Garrott and Eberhardt 1982, Meijer et al. 2011). However, we have not observed a predation event during this study nor during any subsequent years of monitoring (2017-2022) (Roth, unpublished data), suggesting predation risk for pups at dens in our study area is low. Future studies evaluating the relative influence of the possible drivers of predator visits (carrion, feathers/non-food prey remains, hunting foxes) will be beneficial for understanding their attraction to fox dens.

Consistent with our prediction, herbivores, specifically caribou, visited fox dens more often than paired control sites (Fig. 3c). Although caribou may frequent the elevated beach ridges where most dens are located to enhance predator detection or intercept the relatively strong winds that may reduce insect harassment (Hagemoen and Reimers 2002), caribou are more likely to visit fox dens than other areas on the beach ridges. Caribou are likely attracted to the greater biomass and nutrient content of vegetation typically found on dens (Fig. 1, Supplementary Fig. S4; Gharajehdaghipour et al. 2016, Gharajehdaghipour and Roth 2018). Thus, caribou visits to dens are likely driven by a combination of vigilance behavior and enhanced foraging opportunities. For geese, the risk of predation near dens may offset the nutritional benefits of the lush den vegetation, leading to no significant difference in visitation rates between dens and control sites. Due to the size disparity between foxes and caribou, predation risk for caribou is presumably very low at dens, and is likely why we found no difference in herbivore visits between active and inactive dens.

We suggest that fox denning behavior facilitates other wildlife by attracting them to dens through both immediate, direct, and prolonged, indirect pathways. Foxes appear to have an immediate and direct effect on the spatial distribution of predators through their denning activity while occupying dens. Predators visited occupied dens of both Arctic and red foxes, suggesting denning activity by both species may affect predators in a similar manner (although our small sample size precluded effective evaluation of this idea). While these behaviors attract predators in the short-term, they are also the ultimate mechanisms by which fox ecosystem engineering transforms den sites into ecological hotspots. The lush vegetation that grows on Arctic fox dens (Bruun et al. 2005, Gharajehdaghipour et al. 2016, Fafard et al. 2020) continues to indirectly affect the spatial distribution of caribou even if no foxes are present at the dens, demonstrating the long-term effects that fox ecosystem engineering has on other wildlife. Our findings that fox ecosystem engineering affects wildlife differently through these short- and long-term pathways adds to other studies that have demonstrated ecosystem engineering effects may be scale-dependent (Ferry et al. 2020, Johnson-Bice et al. 2022).

Arctic foxes in the southern Arctic are threatened with range contraction in response to the northward expansion of red foxes (Elmhagen et al. 2017, Gallant et al. 2020) and anthropogenic climate change (Verstege 2016). As red foxes expand their presence on the tundra, it remains to be seen whether they will be able to fully replicate the ecosystem engineering role that Arctic foxes play on the tundra. Red foxes tend to leave fewer prey remains scattered around their dens (J.D. Roth, personal observation) and also have smaller average litter sizes than Arctic foxes, suggesting nutrients may be deposited on red fox-occupied dens at lower rates than Arctic fox-occupied dens. As mentioned earlier, the occupation of tundra dens by red foxes is a recent development in our study area, which obscures our ability to evaluate the relative effects each fox species has on other fauna and flora. Continued long-term monitoring of wildlife visits to fox dens may provide more comprehensive evaluations of the non-trophic effects of foxes on other species, and information on how these relationships may vary alongside changes to Arctic and red fox presence on the tundra.

Although most research on predators focuses on trophic interactions (both consumptive and non-consumptive), there is growing recognition that predators impact ecosystem dynamics through important non-trophic pathways. For instance, similar to tundra fox dens, eagle nests are relatively large, long-lasting structures that act as localized biodiversity hotspots in forest environments (Maciorowski et al. 2021). As top predator populations continue to decline throughout many parts of the world (Estes et al. 2011), understanding the complex pathways they impact ecosystems is crucial for advancing the research, conservation, and management of their populations worldwide (Gable et al. 2020). Broadening research on predators to include thorough evaluations of their non-trophic effects provides us with a more enriched view of their ecological role(s) that are integral to functional ecosystems.

## Ethics

This study was conducted under Parks Canada research permit WAP-2013-13539.

## Data accessibility

Data and R code will be uploaded to the Mendeley Data repository upon acceptance.

## Authors’ contributions

S.T.Z. and J.D.R. conceived, designed, and carried out the study. S.M.J-B. prepared and analyzed the data. S.T.Z. and S.M.J-B. wrote the manuscript, with input and revision from J.D.R. All authors gave final approval for publication and agree to be held accountable for the work performed therein.

## Acknowledgements

The CNSC, Wapusk National Park, and Manitoba’s Wildlife Branch provided logistical support. We thank J. Verstege, C. Warret Rodrigues, and students in the University of Manitoba Arctic Field Ecology course in 2015 and 2016 for assistance. J. Verstege, J. Markham, and J. Detwiler provided comments and suggestions on previous drafts.

