## Supplementary material file for "Foxes engineer hotspots of wildlife activity on the nutrient-limited Arctic tundra"

**Supplementary material: details on cataloguing camera events based on animal behavior, and additional details on methods, statistical analysis and results**

**1. Camera trap models and behaviors used to catalog events**

Table S1. List of camera models used in the study.

| Camera model | No. dens* | No. active dens | No. inactive dens |
| --- | --- | --- | --- |
| Bushnell Trophy Cam 119466 | 8 | 1 | 7 |
| Moultrie Game Spy C-90i | 1 | 1 | 0 |
| Reconyx PC800 | 7 | 4 | 3 |

*for each den, another camera of the same model was used at the control site

The behavior of animals was used to determine whether images of animals collected by camera traps should be counted as a visit to an area (den or control). Behaviors were described separately for mammals and birds and were prioritized into three categories reflecting the strength of evidence of a visit (Table S2). An animal exhibiting any behaviors of category I was included as a visit, regardless of the duration of the visit (Figs S1, S2, and Fig. 2 from the main text). An animal exhibiting a behavior in category II was only considered to have visited the area if the length of time the animal was photographed was >1 min. An animal exhibiting only category III behaviors was not included as a visit.

Table S2. Description of behavior of animals used for determining wildlife visits. Behavioral categories (I, II, III) reflect the priority of use for describing a visit to an area. Any Category I behavior was counted as a visit, whereas Category II counted only if the duration exceeded 1 min, and an animal exhibiting only Category III behaviors was not included as a visit. Some behaviors were described separately for mammals (bears, wolves, caribou and hares) and birds (eagles, ravens, owls, cranes, and geese).

| **Behavior** | **Description** |
| --- | --- |
| **Category I** | |
| Foraging | *Mammal*: head at or below the knee level and the muzzle held no higher than the knee when animal is in the grass; or the muzzle depressed to its ankle level when animal is not surrounded by grass. *Note: the knee and ankle height are measured on the leg that contacts the ground. |
|  | *Bird*: neck depressed, head lower than the body plane and eyes looking down at the ground; the bill may or may not be touching the ground. |
| Resting | *Mammal*: legs retracted underneath the body or head and legs outstretched while lying on the ground. |
|  | *Bird*: Both legs retracted underneath the body. Head and neck either held up or retracted to chest region. |
| Digging | *Mammal*: using forelimbs (single or both) to remove objects (dirt, vegetation etc.) from ground surface or through the ground. |
| Interspecific agonistic | Individuals of different species stand still and look at one another at minimum of five meters distance or individual of one species is running after another individual of a different species. |
| Urinating | *Mammal*: standing on three legs with one hind leg elevated in the air or slightly extending hind legs and urine comes out of the urethra area. |
| Nursing | *Mammal*: calf has its head slightly elevated and close to the belly of the female, interpreted as suckling from the female. |
| Landing | *Bird*: descending from flight and extending legs to approach the ground, suggesting a transition of behaviors from flying to other activities in the area. |
| **Category II** | |
| Locomotion | Moving legs alternatively while keeping the body plane roughly parallel to the ground (e.g. walking or running). |
| Standing | In a quadrupedal or bipedal stance without leg movement. |
| **Category III** | |
| Camera | Any of the following: |
| interaction | a) *Mammal*: standing by the camera with body partially captured by the camera; |
|  | b) *Mammal*: approaches and makes contact with the camera by sniffing with nose or chewing with mouth or rubbing with part of the body; |
|  | c) *Bird*: sitting or standing on top of the camera and having either claws or tail feathers captured by the camera. |
| Flying | Any of the following: |
|  | a) *Bird*: standing on the ground while extending wings to start the upward and downward strikes; |
|  | b) *Bird*: in the air with wings spread. |

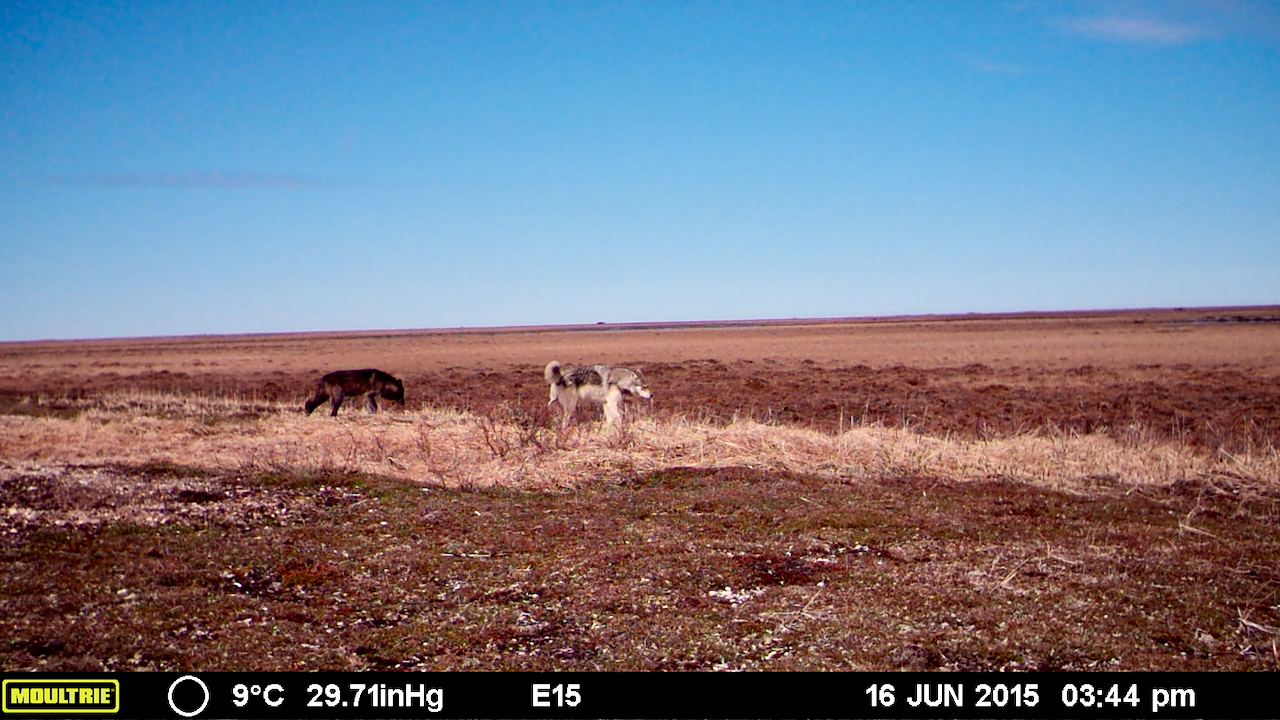

Figure S1. Two wolves visiting a fox den, with one urinating on the den site.

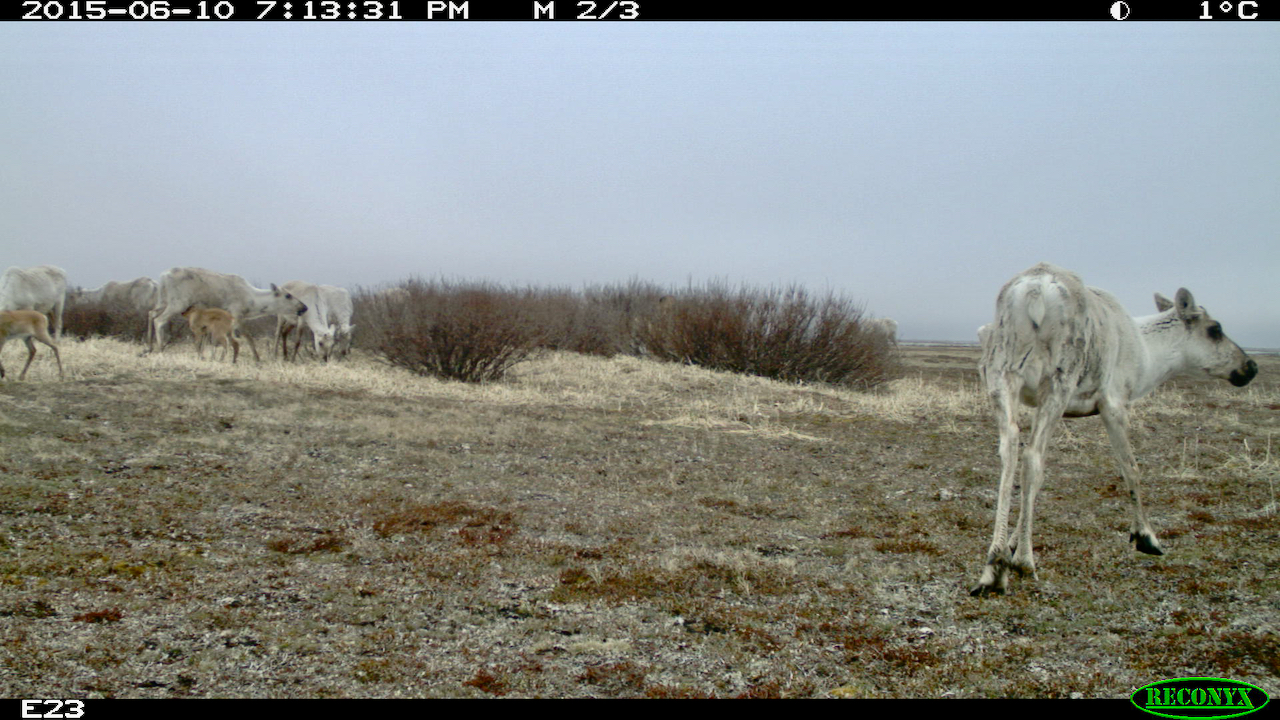

Figure S2. Caribou foraging and nursing at a den site in early June.

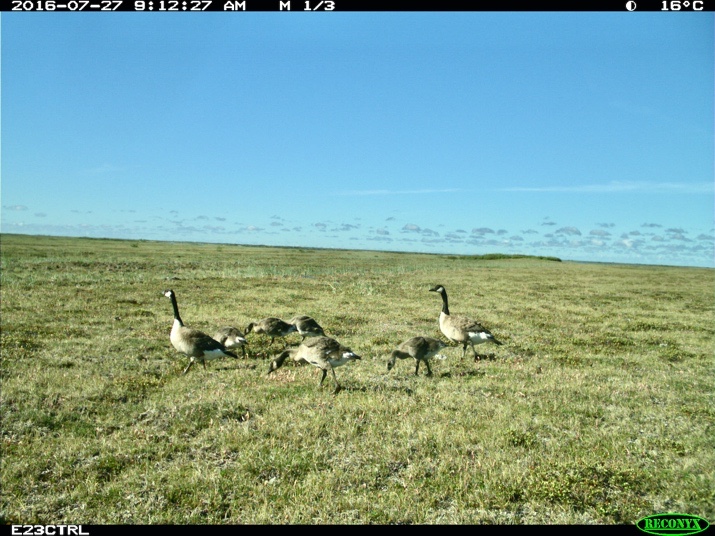

Figure S3. Canada geese foraging at a control site.

**2. Full modeling results comparing wildlife visits to dens/control areas, and active/inactive dens**

Table S3. Generalized linear mixed model results comparing wildlife captures between den and control sites (SE = Standard error; CI = confidence intervals).

| Comparison | Distribution | Parameter | Estimate | SE | 95% CI | Z-value | *p*-value |
| --- | --- | --- | --- | --- | --- | --- | --- |
| Total wildlife captures | negative binomial | Intercept | –1.684 | 0.46 | (–2.586, –0.782) | –3.660 | <0.001 |
|  |  | Treatment_Den_ | 0.994 | 0.33 | (0.350, 1.638) | 3.026 | 0.002 |
|  |  | Year_2016_ | –0.021 | 0.43 | (–0.861, 0.820) | –0.048 | 0.962 |
| All herbivores | negative binomial | Intercept | –1.826 | 0.52 | (–2.854, –0.798) | –3.483 | <0.001 |
|  |  | Treatment_Den_ | 0.830 | 0.35 | (0.146, 1.515) | 2.377 | 0.017 |
|  |  | Year_2016_ | 0.073 | 0.48 | (–0.860, 1.007) | 0.154 | 0.878 |
| All predators | Poisson | Intercept | –7.409 | 1.18 | (–9.714, –5.103) | –6.299 | <0.001 |
|  |  | Treatment_Den_ | 3.638 | 1.01 | (1.652, 5.623) | 3.591 | <0.001 |
|  |  | Year_2016_ | –0.494 | 0.44 | (–1.351, 0.363) | –1.129 | 0.259 |
| Species richness | Poisson | Intercept | –3.460 | 0.29 | (–4.025, –2.895) | –12.009 | <0.001 |
|  |  | Treatment_Den_ | 0.647 | 0.26 | (0.131, 1.162) | 2.457 | 0.014 |
|  |  | Year_2016_ | –0.297 | 0.29 | (–0.864, 0.271) | –1.024 | 0.306 |
| *Post-hoc models* | |  |  |  |  |  |  |
| Caribou | negative binomial | Intercept | –2.453 | 0.51 | (–3.448 –1.459) | –4.834 | <0.001 |
|  |  | Treatment_Den_ | 0.850 | 0.37 | (0.131, 1.569) | 2.316 | 0.021 |
|  |  | Year_2016_ | 0.068 | 0.46 | (–0.837, 0.972) | 0.146 | 0.884 |
| Geese* | negative binomial | Intercept | –2.433 | 1.75 | (–5.854, 0.988) | –1.394 | 0.163 |
|  |  | Treatment_Den_ | 0.743 | 1.33 | (–1.861, 3.346) | 0.559 | 0.576 |
|  |  | Year_2016_ | –0.415 | 1.60 | (–3.552, 2.723) | –0.259 | 0.796 |
| *Includes all geese species | | | | | | | |

Table S4. Generalized linear mixed model results comparing wildlife captures between active and inactive dens (SE = Standard error; CI = confidence intervals).

| Comparison | Distribution | Parameter | Estimate | SE | 95% CI | Z-value | P-value |
| --- | --- | --- | --- | --- | --- | --- | --- |
| All predators | Poisson | Intercept | –1.494 | 0.402 | (–2.283, –0.706) | –3.716 | <0.001 |
|  |  | Activity status_Inactive_ | –2.556 | 0.525 | (–3.584, –1.527) | –4.871 | <0.001 |
|  |  | Year_2016_ | –1.187 | 0.406 | (–1.982, –0.392) | –2.925 | 0.003 |
| All herbivores | negative binomial | Intercept | 0.190 | 0.563 | (–0.914, 1.295) | 0.338 | 0.736 |
|  |  | Activity status_Inactive_ | –0.474 | 0.657 | (–1.762, 0.815) | –0.720 | 0.471 |
|  |  | Year_2016_ | –0.857 | 0.626 | (–2.084, 0.370) | –1.369 | 0.171 |
| Species richness | Poisson | Intercept | –2.428 | 0.185 | (–2.791, –2.066) | –13.133 | <0.001 |
|  |  | Activity status_Inactive_ | –0.718 | 0.215 | (–1.139, –0.296) | –3.334 | <0.001 |
|  |  | Year_2016_ | –0.283 | 0.221 | (–0.716, 0.149) | –1.285 | 0.199 |

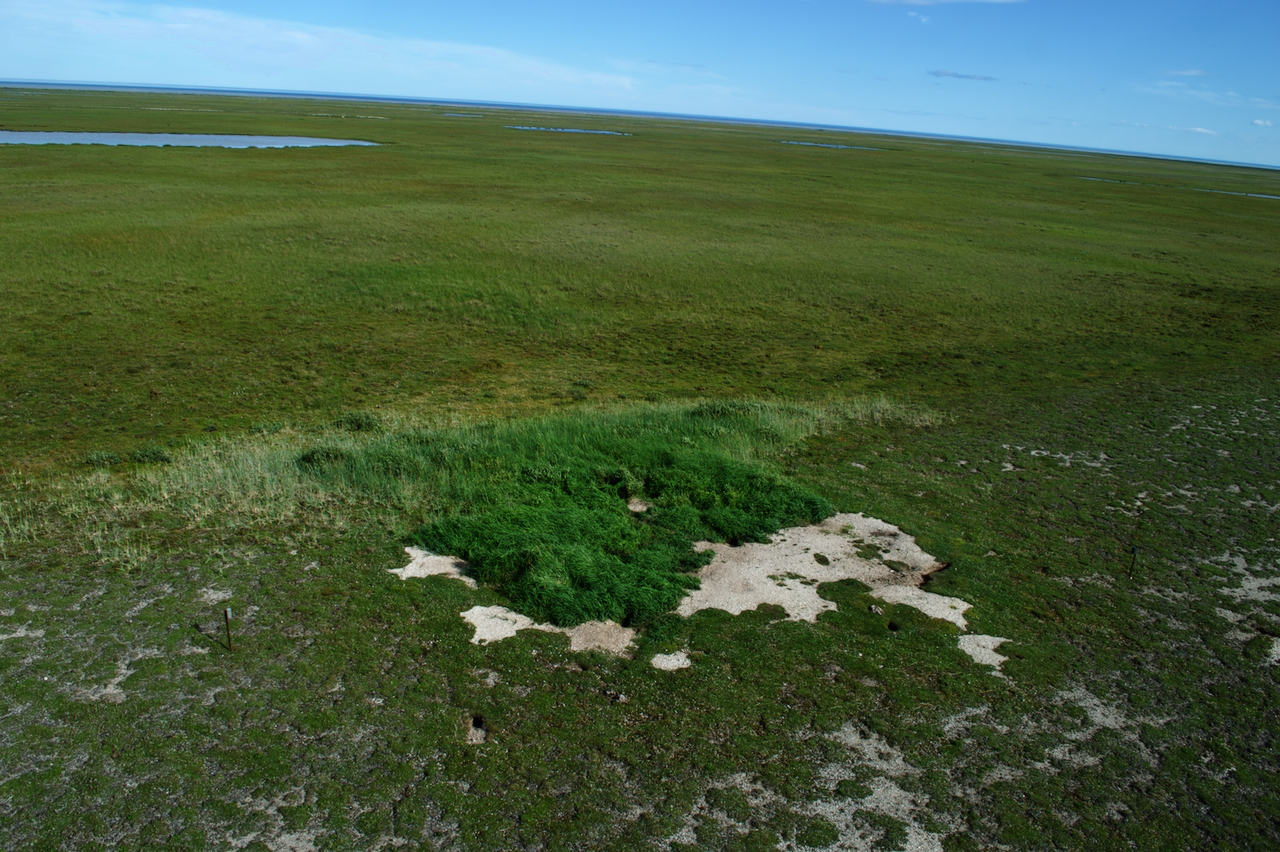

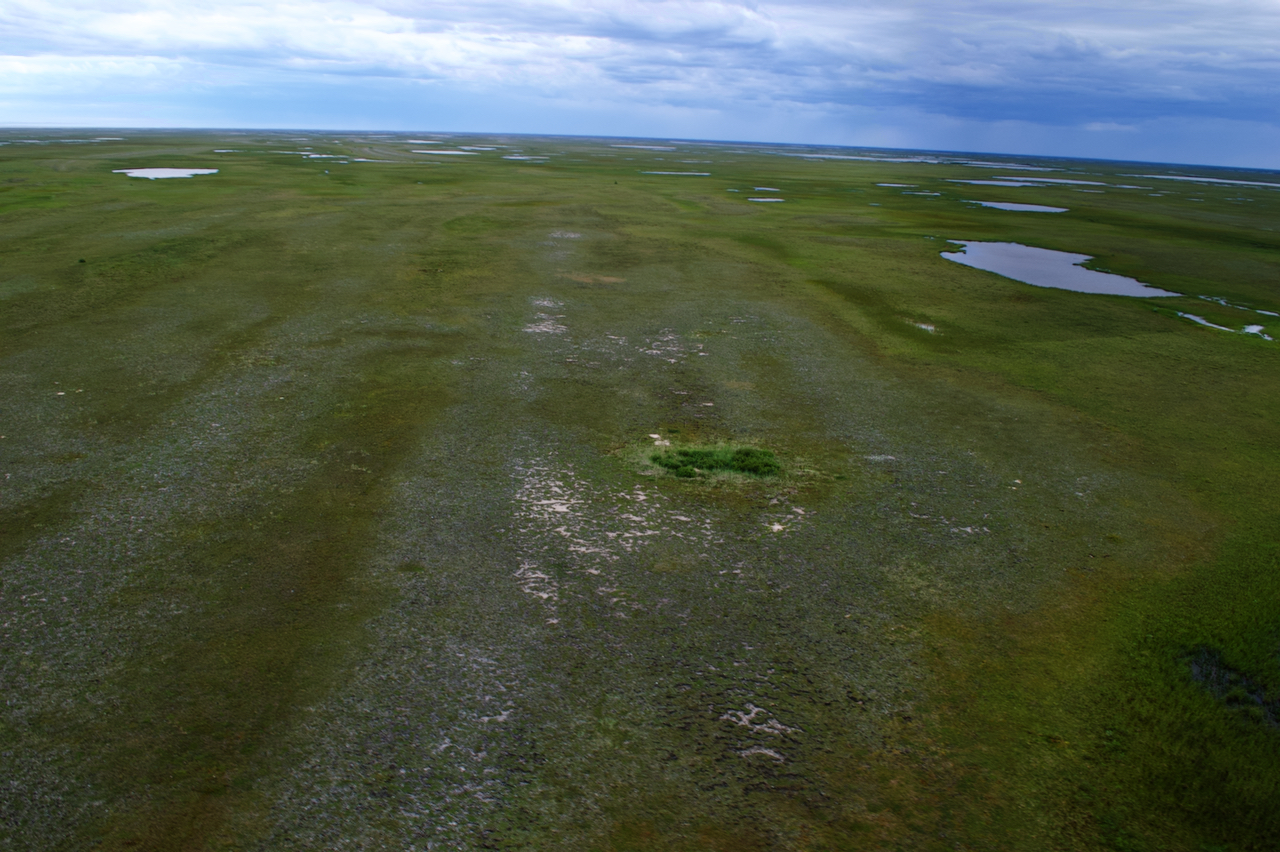

Figure S4. Aerial photos (top and bottom) of Arctic fox dens in Wapusk National Park taken in August, 2020. The enriched, tall vegetation on the den sites is distinctive on the tundra landscape. Nonetheless, the bottom photo also clearly shows the relative homogeneity of the beach ridges in Wapusk.
